# ERK activity as a key player in determining cardiac cell fate choices

**DOI:** 10.64898/2026.02.18.706664

**Authors:** Karin Farkas, Elisabetta Ferretti

## Abstract

Heart development is supported by diverse cell types originating from the lateral plate mesoderm (LPM), including the first heart field (FHF), the second heart field (SHF), which is a part of pharyngeal mesoderm, and the newly described juxta-cardiac field (JCF) that harbors progenitor cells of the epicardium. While FGF-MEK-ERK signalling has been implicated in various developmental mechanisms, its role in cardiac specification remains elusive. This signalling pathway involves autoinhibitory loops, acting at translational and post-translational levels, resulting in pulses of ERK activity. We hypothesized that this alternating ERK activity could direct binary cell fate choices during mesodermal specification. Using an *in vitro* system, we found that inhibition of ERK activity in the LPM, before cardiac commitment, resulted in enriched gene expression of JCF and pharyngeal mesoderm/SHF markers, with high proepicardial marker levels, at the expense of FHF-like cardiomyocyte markers. Our findings reveal a novel contribution of ERK signalling in cell differentiation within the cardiac lineages.

## Introduction

Cardiovascular diseases are a major cause of global mortality, highlighting the need for a better understanding of heart development and disease [1]. Human pluripotent stem cells (PSCs) provide a valuable tool for studying development and diseases, as they can be differentiated into various cell types [2–5]. The use of PSCs to model development and diseases and the progress in regenerative treatment strategies utilizing stem cells show great promise for the future of cardiac medicine [6, 7]. Understanding the complex cellular and molecular interactions regulating heart development is critical for modelling and treating cardiac diseases.

During development, the heart is formed by multiple cell sources in a spatially and timely coordinated manner [8–10]. A large portion of heart progenitor cells stem from the mesoderm, and the specification of the cardiac precursors takes place when the cells migrate out of the primitive streak (PS) [8, 10]. In the third gestational week, a cell population called the FHF forms the cardiac crescent, which acts as a scaffold as the heart grows [8–10]. The FHF mainly contributes to the future left ventricle, whereas the cell population joining from the pharyngeal mesoderm, the SHF, contributes to the right ventricle, the two atria and the outflow tract (OFT) [8–11]. The cardiac neural crest is of ectodermal origin and forms the OFT, the atrioventricular septum, valves and ganglia [8–10]. Another extra-cardiac cell source is proepicardium, a cell cluster that later migrates on the surface of the heart as epicardium [12, 13]. Surrounding the heart, the epicardium acts as a signalling center ensuring proper cardiac growth and patterning, and differentiates into the non-cardiomyocyte lineages of the heart, such as coronary vessels and cardiac fibroblasts [14, 15].

During early cardiac specification events, mesodermal progenitors are exposed to agonists and antagonists of BMP4, FGF and WNT from the surrounding tissues [9, 10]. The FGF signalling pathway has been reported to be crucial during development in diverse contexts [16–18]. Upon binding to the receptors, FGF can activate several intracellular pathways, including the MEK/ERK pathway [16–18]. This pathway is mediated by the kinases RAF-MEK-ERK cascade, in which rapid signal amplification is achieved by sequential phosphorylation of these components [19]. Finally, phosphorylated ERK (pERK) activates various transcription factors, leading to gene expression changes [16, 20]. The pathway also involves autoinhibitory loops that can be mediated either by (de)phosphorylation or by translation of inhibitors such as DUSP or Sprouty proteins [16, 21]. These negative feedback loops have been shown to generate pulsing ERK activity patterns, creating intrinsic clocks dictating the execution of developmental programs and directing cell fate commitment [22–24]. For example, pulsing ERK activity is crucial for generating periodic segments in presomitic mesoderm [24].

In this study, we hypothesized that temporal modulation of ERK activity could direct cell fate acquisition in the cardiac lineages. We inhibited the ERK pathway at the LPM stage prior to cardiac commitment. We found that a transient inhibition resulted in alteration of cell type markers from FHF to a wider progenitor pool of cardiac and extra-cardiac lineages including the proepicardium. While the derivation of proepicardium from PSCs has been previously achieved by timely modulation of the WNT signalling [25–29], the current study shows that temporal MEK/ERK inhibition can similarly induce proepicardium formation. Our findings add further layers to the understanding of cardiac specification mechanisms, provide a faster protocol for deriving proepicardium, which is implicated in cardiac regeneration [14, 15], and could have crucial implications for research in stem cell-based treatments.

## Results

### MEK/ERK inhibition in LPM induces distinct splanchnic mesoderm genes

To evaluate the effect of MEK/ERK inhibition during cardiac induction, we first established a controllable differentiation protocol. To this end, we improved a stepwise mesoderm differentiation protocol for human embryonic stem cells (ESCs) with a distinct set of cytokines administered on each day [30]. We adapted the protocol to a monolayer culture and to chemically fully defined conditions to minimize variability (Fig. 1A). In the control condition, a continuous mesh-like cardiomyocyte layer with a wave-like beating pattern could be observed on day 8 (Fig. 1B, Video S1-2). After reaching the LPM stage, these control cells were treated with the cardiac mesoderm medium containing FGF2. At this stage, we applied a potent MEK inhibitor PD0325901 (PD) instead of FGF2 for 24 h, to achieve an effective interference with the FGF-MEK-ERK pathway. PD allosterically binds to MEK and inhibits its kinase activity, preventing ERK phosphorylation and activation [31]. The PD-treated cells under this condition resulted in isolated twitching on day 8 (Fig. 1B, Video S3-4). These findings suggest that a transient PD treatment, or absence of FGF2 in the medium, had a lasting effect observed in the morphology of the beating structures. This variability suggested potential differences in transcriptional profiles, that we investigated next.

**Figure 1.**
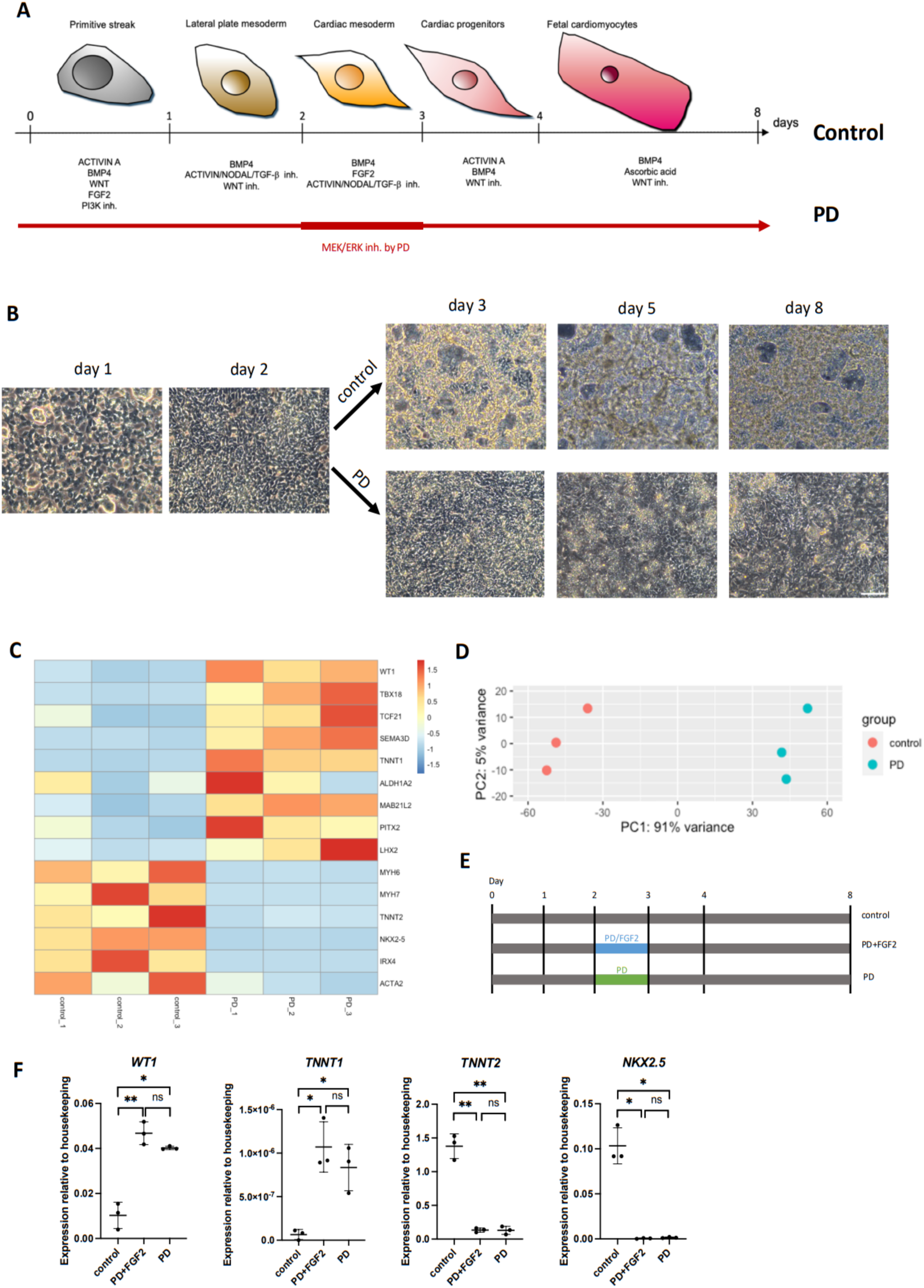
Transient MEK/ERK inhibition sustainably alters transcriptomics. (**A**) Schematic representation of the stepwise cardiac differentiation protocol. In the PD-treated conditions, MEK/ERK is inhibited by PD on day 3 for 24 h. (**B**) Brightfield images at 20x magnification showing representative cell morphology in control and PD-treated cultures. The control culture gradually forms a mesh-like cell network with wave-like beating, whereas PD-treated cells develop into a monolayer with isolated twitching. Scale bar, 100 μm. (**C**) Heatmap showing expression patterns of selected cardiogenesis-associated genes (epicardial, pharyngeal and cardiomyocyte genes) of control and PD-treated 2D cultures on day 8 from bulk RNAseq experiment. Proepicardial genes *WT1, TBX18, TCF21* and *SEMA3D* as well as pharyngeal genes *PITX2* and *LHX2* are upregulated in the PD-treated cells. Results were obtained from three biological replicates. Scale, Z-score. (**D**) PCA plot of the bulk RNAseq experiment indicating a clear separation of control and PD-treated samples. (**E**) Experimental setup to test the effect of PD in the presence and absence of FGF2 in our protocol. Cells were collected on day 8 and analyzed by RT-qPCR. (**F**) RT-qPCR results showing that PD treatment promotes proepicardial markers in presence and absence of FGF2. Error bars indicate means with standard deviations. Three biological replicates were analyzed in two technical replicates. Not significant (ns) P > 0.05, * P ≤ 0.05, ** P ≤ 0.01 (t-test).

To this end, we collected the control and PD-treated cells on day 8 when the cultures clearly showed beating structures, and subjected the samples to bulk RNA sequencing (RNAseq) (Fig. 1C). Principal component analysis (PCA) showed distinct RNA expression profiles for control and PD conditions (Fig. 1D). Next, we compared the expression levels of cardiogenesis-associated genes in both conditions. The cells subjected to PD treatment showed upregulation of genes associated with proepicardium, including *WT1*, *TBX18*, *TCF21* and *SEMA3D* [32–36], as well as the troponin *TNNT1* which has been reported to be enriched in epicardium [37, 38]. However, *ALDH1A2*, the marker associated with mature epicardium [29, 39], was not significantly upregulated, suggesting that these cells represent the progenitor state. In addition, the marker of the recently identified JCF, *MAB21L2*, was also elevated in PD-treated cells, consistent with the finding that JCF harbors proepicardial progenitors [40–42]. Of note, the pharyngeal mesoderm markers *PITX2* and *LHX2* were also elevated in PD-treated cells [43, 44]. In contrast, cardiomyocyte, FHF and smooth muscle markers, including *MYH6, MYH7, TNNT2, NKX2.5, IRX4* and *ACTA2* [45–48], were upregulated in the control cells. Thus, the control cells could represent FHF-like cardiomyocytes, whereas PD-treated cultures could correspond to the cells from a wider field that can contribute to the proepicardium and pharyngeal mesoderm including the SHF. However, the distinction between proepicardium and septum transversum mesenchyme (STM) is challenging due to the lack of specific markers [28, 49–51], hindering a more precise cell type determination.

In our protocol, FGF2 was omitted from the cardiac mesoderm medium when supplying PD, to prevent the overactivation of signalling pathways diverting from FGF2, such as JNK, PI3K/AKT and JAK/STAT [52]. To investigate if the MEK/ERK inhibition, rather than the reduction of the feed into other FGF-associated pathways, was responsible for the observed differences upon PD treatment, we evaluated the gene expression patterns of three distinct conditions on day 8: control, PD+FGF2 and PD (Fig. 1E). Quantitative PCR analysis indicated that the expression patterns of cardiac genes were similar between PD+FGF2 and PD conditions and differed from the control. The expression of (pro)epicardial markers *WT1* and *TNNT1* were upregulated while the cardiomyocyte markers *TNNT2* and *NKX2.5* were downregulated in both PD+FGF2 and PD conditions (Fig. 1F). These findings suggest that the observed gene expression changes were due to MEK/ERK inhibition rather than the attenuation of the signalling pathways diverting from FGF.

To sum up, these results imply that the MEK/ERK inhibition at the LPM stage directs the transcriptomic signatures towards non-FHF lineages within splanchnic mesoderm, such as proepicardium and SHF. Previous studies suggested that cardiomyocytes, proepicardium, SHF and STM originate from LPM [53–56], which is consistent with our results.

### Proepicardial markers are observed upon MEK/ERK inhibition in LPM

Given that an elevated ERK activity was expected after alleviating its inhibition, we analyzed the scope and duration of MEK/ERK inhibition by PD in our experimental settings (Fig. 2A). To this end, we collected cell lysates from day 2 just before PD administration, 6 h into PD treatment, on day 3, 4 and 8, and assessed the levels of total and phosphorylated MEK and ERK by western blotting (Fig. 2B, S1). An accumulating trend of phosphorylated, active MEK (pMEK) and vanishing trend of pERK were observed already after 6 h of PD treatment. The decreased trend in pERK levels compared to the control lasted at least until day 4, i.e. 1 day after PD removal. These results imply that the effect of PD on the ERK activity persists even after PD removal and the pERK levels in PD-treated cells do not increase compared to the control at the tested time points.

**Figure 2.**
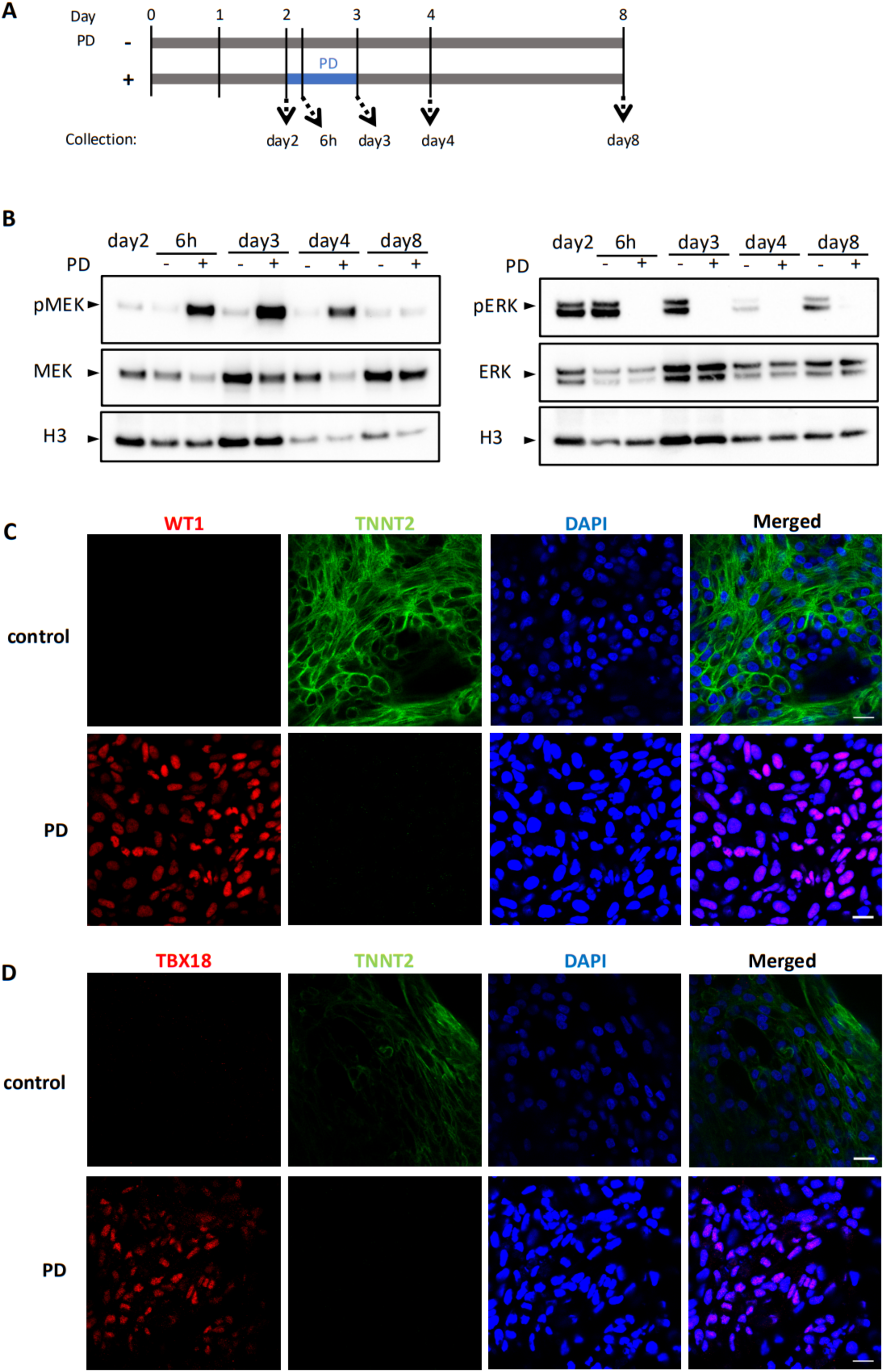
Proepicardial markers are expressed upon MEK/ERK inhibition at the LPM stage. (**A**) Schematic representation of the western blot experiment assessing MEK and ERK phosphorylation status in control cells and upon PD treatment. Cell lysates were collected on day 2 before PD administration, 6 h into treatment, day 3, 4 and 8 to follow the phosphorylation patterns over time. (**B**) Western blotting of phosphorylated and total MEK and ERK. Due to the inhibition by PD, pMEK accumulates, whereas the downstream pERK decreases. Two biological replicate experiments were performed. (**C-D**) Immunofluorescence of 2D cultures on day 8 demonstrating the cardiomyocyte marker TNNT2 in the control and proepicardial markers WT1 and TBX18 in the PD-treated cells. Two biological replicate experiments were produced and three fields were randomly imaged per condition per experiment. Representative images are shown. Scale bars, 20 μm.

To further consolidate the proepicardial identity of PD-treated cells, we analyzed the levels of proepicardial and muscle markers on day 8 by immunofluorescence (Fig. 2C-D). PD-treated cells exhibited WT1 and TBX18 but almost no TNNT2, whereas control cells showed TNNT2 but almost no WT1 or TBX18. These markers were expressed in a non-uniform manner, in agreement with the heterogeneity of the cell populations found in PSC-derived cardiac cultures [57]. Overall, these results support the RNAseq findings, indicating that PD-treated cells express proepicardial markers.

### MEK/ERK inhibition promotes proepicardial gene expression in organoid cultures

In the control condition, cells often spontaneously formed rounded 3D structures, implying that the aggregate architecture more closely mimics the physiological environment during cardiac differentiation compared to the monolayer. Thus, we envisaged that the cellular 3D architecture is relevant during cardiogenesis, and investigated the effect of the MEK/ERK inhibition during cardiac differentiation using a 3D organoid culture. To this end, we generated cardiac organoids starting from embryoid bodies with the same cytokines employed in the monolayer differentiation protocol (Fig. 3A-B, Video S5-8). We performed bulk RNAseq on day 8 organoid samples to analyze the expression profiles of cardiogenesis-associated genes (Fig. 3C). As in the 2D monolayer cultures, the proepicardial markers *WT1, TBX18, TCF21, SEMA3D* and *TNNT1* were upregulated in PD-treated cells, whereas muscle markers *MYH6, MYH7, NKX2.5, IRX4* and *ACTA2* were downregulated. The JCF marker *MAB21L2* was also upregulated in PD-treated cells, with a non-significant increase of the pharyngeal markers *PITX2* and *LHX2*, although a trend was noticed.

**Figure 3.**
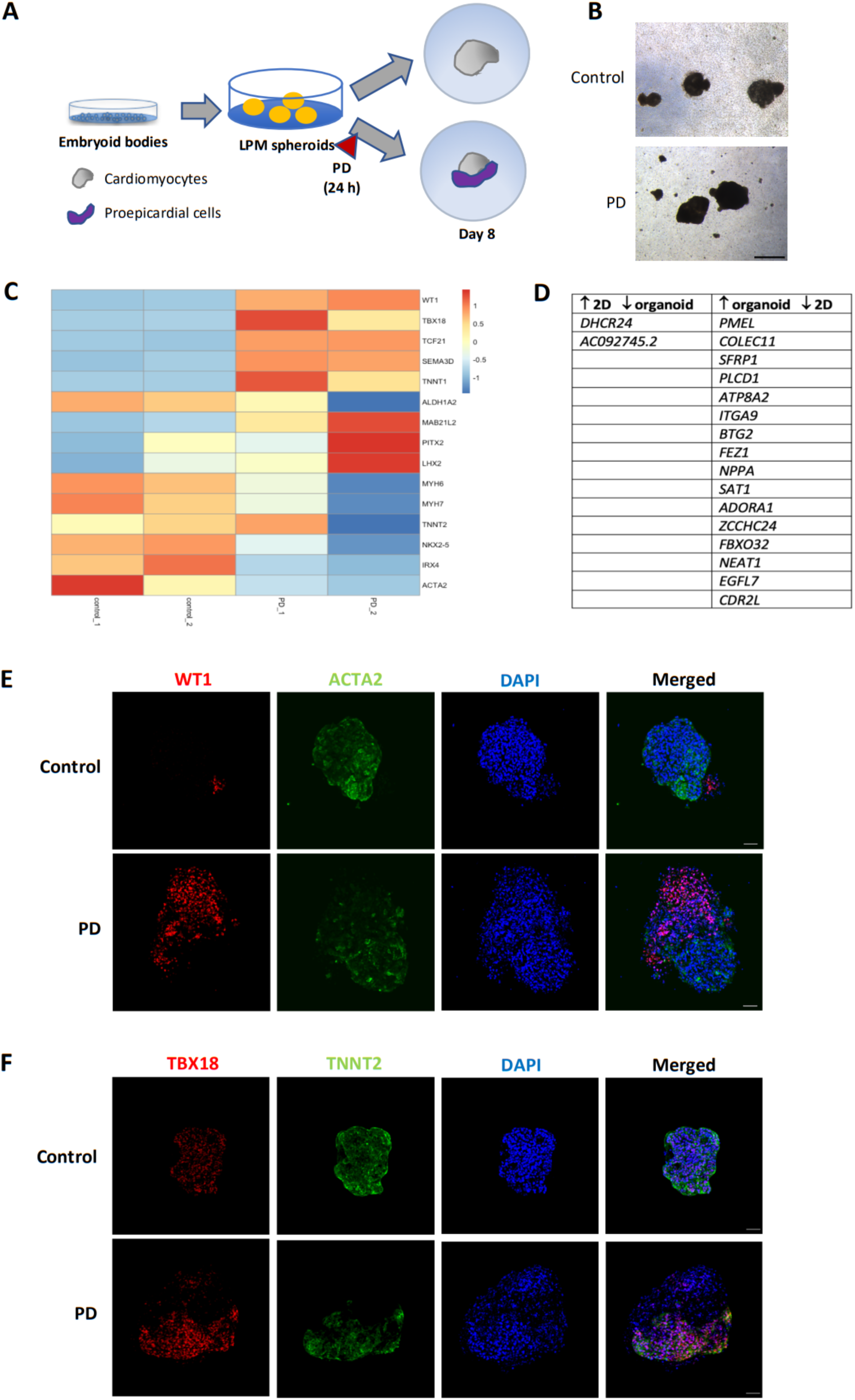
MEK/ERK inhibition induces proepicardial markers in the 3D organoid culture. (**A**) Schematic of the cardiac organoid differentiation. Embryoid bodies were differentiated with the same cytokine cocktails used in the monolayer culture. (**B**) Representative brightfield images of control and PD-treated day 8 organoids at 5x magnification. Scale bar, 500 μm. (**C**) Heatmap of the expression levels of selected cardiogenesis-associated genes (epicardial, pharyngeal and cardiomyocyte genes) in day 8 organoid cultures assessed by bulk RNAseq. As in the 2D cultures, proepicardial genes are upregulated and cardiomyocyte genes are downregulated in the PD-treated cells compared to the control. Two biological replicate experiments were performed. Scale, Z-score. (**D**) Assessment of the effect of the cell architecture on PD-treated cells based on bulk RNAseq experiments. Some genes show opposite trends in 2D versus 3D cultures upon PD treatment. (**E-F**) Immunofluorescence of day 8 control and PD-treated organoids demonstrating muscle (ACTA2, TNNT2) and proepicardial markers (WT1, TBX18). Two biological replicate experiments were performed and three random organoids were imaged per experiment per condition. Representative images are shown. Scale bars, 50 μm.

Thus, during 3D organoid differentiation, MEK/ERK inhibition at the LPM stage mainly elevates the expression levels of proepicardial markers at the expense of muscular markers.

Next, we addressed the transcriptomic differences between the 2D and 3D cultures upon PD treatment (Fig. 3D). The RNAseq analysis revealed that *DHCR24*, encoding the final enzyme in the cholesterol biosynthesis pathway and involved in protecting against cardiomyocyte apoptosis [58, 59], was upregulated in the PD-treated 2D culture but downregulated in the 3D culture. Conversely, the expression levels of 16 genes were downregulated in the 2D culture and upregulated in the 3D culture in PD-treated conditions. Several of these genes have anti-proliferative or anti-migratory roles, including *SFRP1, PLCD1, BTG2, ATP8A2, ITGA9, FBXO32* and *NPPA* [60–67]. *SFRP1,* highly expressed in the developing heart, antagonizes vascular cell proliferation by inhibiting the WNT signalling [60]. Likewise, PLCD1 and BTG2 have also been reported to inhibit the WNT pathway and to suppress proliferation, invasion and migration of cancer cells [61, 62]. ATP8A2, ITGA9 and FBXO32 have been shown to suppress proliferation and/or migration in disease models [63–65]. Notably, *NPPA* has been associated with cardiogenesis, where it is expressed in the atrial and ventricular myocardium during development, and has also been shown to inhibit proliferation of vascular smooth muscle cells and endothelial cell growth [66, 67]. Furthermore, *Nppa* expression levels and WNT activity have been found to be inversely correlated in a mouse cardiomyopathy model [68], implying that NPPA might negatively regulate WNT signalling. Overall, the activation of genes that suppress proliferation and/or migration in the PD-treated organoids implies some differentially activated pathways in 2D and 3D cultures upon PD treatment.

Next, we employed immunofluorescence to investigate the expression of proepicardial and muscle markers in 3D organoid cultures at the protein level (Fig. 3E-F). As in the 2D cultures, the cell populations in day 8 organoids were non-uniform. Myocyte markers ACTA2 and TNNT2 as well as proepicardial markers WT1 and TBX18 were found in both the control and PD-treated organoids. Thus, the effect of PD treatment may not be as profound as in the 2D environment. Overall, the results suggest that the PD treatment increases the proepicardial marker expression in the organoids at least at the transcriptomics level.

### Single-cell RNAseq reveals gene expression patterns in early progenitor cells

Given the observed cell heterogeneity, we addressed the composition of the cell populations during cardiac induction to understand the differentiation dynamics from LPM. To this end, we conducted single-cell RNAseq (scRNAseq) analysis on cryopreserved cells from control and PD-treated 2D cultures at three time points, day 2 (LPM), day 3 and day 8 to address the gene expression changes over time. By Uniform Manifold Approximation and Projection (UMAP), we found that the five sequenced samples largely delineated the five cell groups (Fig. 4A, S2). Next, we focused on the genes regulating the ERK pathway to understand how the PD treatment affected the ERK signalling over time (Fig. 4B). On day 3, control cells were enriched in the ERK suppressor *DUSP6*, while PD-treated cells exhibited higher *FGFR2* levels. On day 8, *DUSP6* and *FGFR2* enrichment in the control and PD-treated cells, respectively, persisted, suggesting a long-term impact of a transient MEK/ERK inhibition.

**Figure 4.**
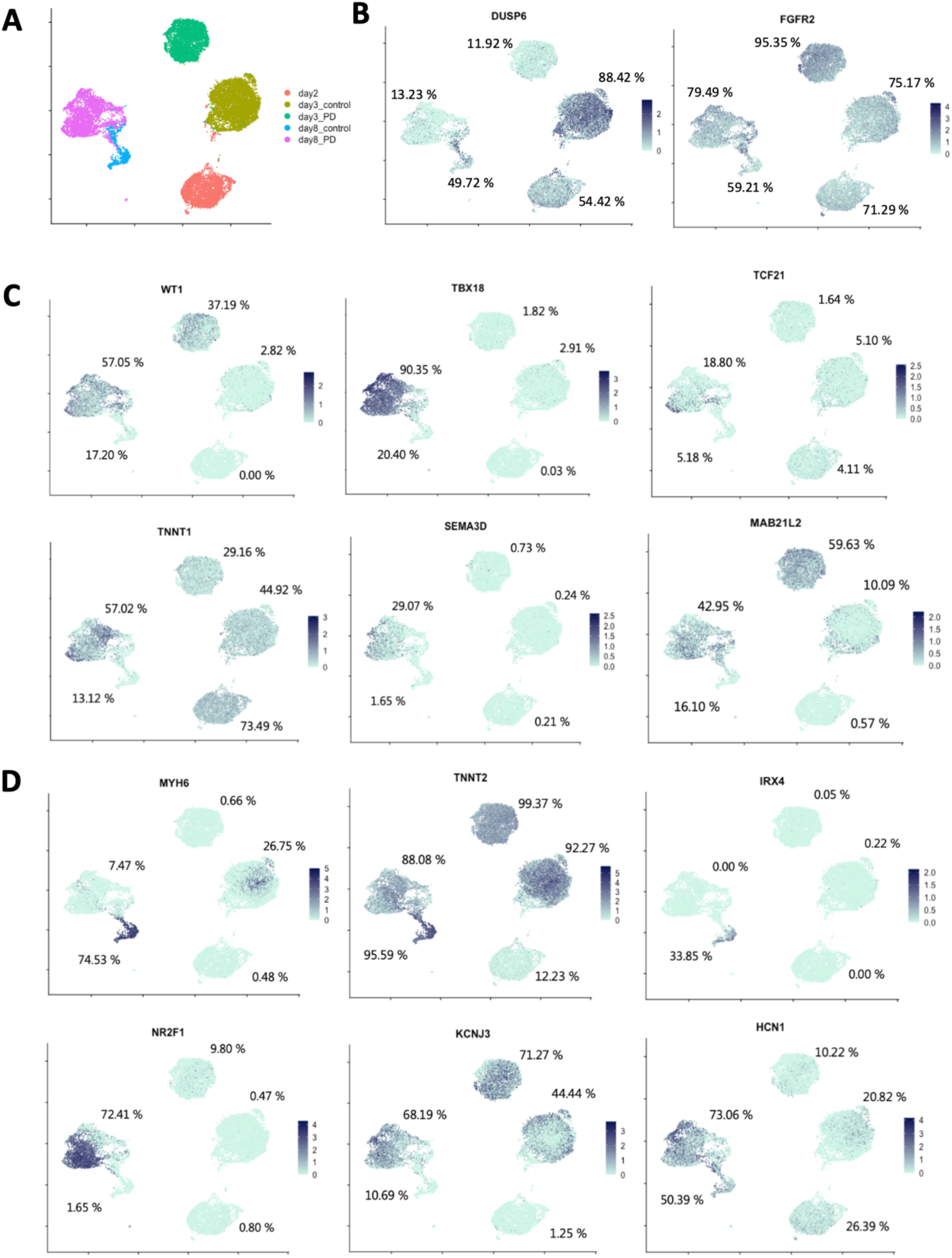
Single-cell RNAseq reveals expression patterns of proepicardial and myocardial genes over time. (**A**) The five sequenced samples from day 2, day 3 and day 8 are separated from each other on the UMAP plot. (**B**) UMAP plots showing differential regulation of genes associated with the FGF-ERK signalling pathway in control and PD-treated cells. (**C**) UMAP plots demonstrating the upregulation of proepicardial genes in day 3 and/or day 8 PD-treated cultures. (**D**) UMAP plots showing the differential expression patterns of muscle and pacemaker genes in control and PD-treated cells. SHF-associated genes *NR2F1* and *KCNJ3* are observed in day 8 PD-treated cells, consistent with their pharyngeal origin. The indicated percentage refers to the fraction of the cells whose normalized transcript count is higher than 1.5, which corresponds to 0.41 on the shown natural log-transformed scale.

Expectedly, PD treatment led to the elevated expression of proepicardial markers *WT1*, *TBX18*, *TCF21, TNNT1* and *SEMA3D* on day 8 (Fig. 4C). We also observed the expression of early proepicardial marker *WT1* in a fraction of day 3 PD-treated cells, consistent with its role in common progenitors of proepicardium, kidney and liver [69–71]. Additionally, we found the expression of the JCF marker *MAB21L2* in both day 3 and day 8 PD-treated cells, consistent with *MAB21L2* representing an early marker of STM [40, 72]. Focusing on cardiomyocyte markers, many genes such as *MYH6, TNNT2* and *IRX4* showed elevated expression levels in day 8 control cells compared to day 8 PD-treated cells, as expected (Fig. 4D). However, some atria-specific genes, such as *NR2F1* and *KCNJ3* [73], were enriched in PD-treated cells (Fig. 4D), supporting their pharyngeal/SHF identity. Moreover, the pacemaker marker *HCN1* [74, 75] was found to be enriched in day 8 PD-treated cells (Fig. 4D). Of note, *TBX18* has also been implicated in the induction of the pacemaker identity [76, 77], suggesting that a subset of PD-treated cells might represent pacemaker progenitor cells. Thus, the PD-treated cell population could contain several progenitor cell types or the common progenitors of several lineages, including proepicardial, SHF and pacemaker precursors.

### PD-treated cells resemble *in vivo* epicardium

Subsequently, to confirm the identity of the PD-treated cells and to evaluate their representation of *in vivo* (pro)epicardium, we compared gene expression profiles between the PD-treated cells and the existing scRNAseq data of fetal epicardium [38] (Fig. 5A). Integration of transcriptomics data of cells annotated as epicardium with the day 8 PD-treated cells showed comparable expression levels of epicardial marker genes in both populations (Fig. 5B-C, S3), indicating that the day 8 PD-treated cells are a preeminent representation of *in vivo* epicardium.

**Figure 5.**
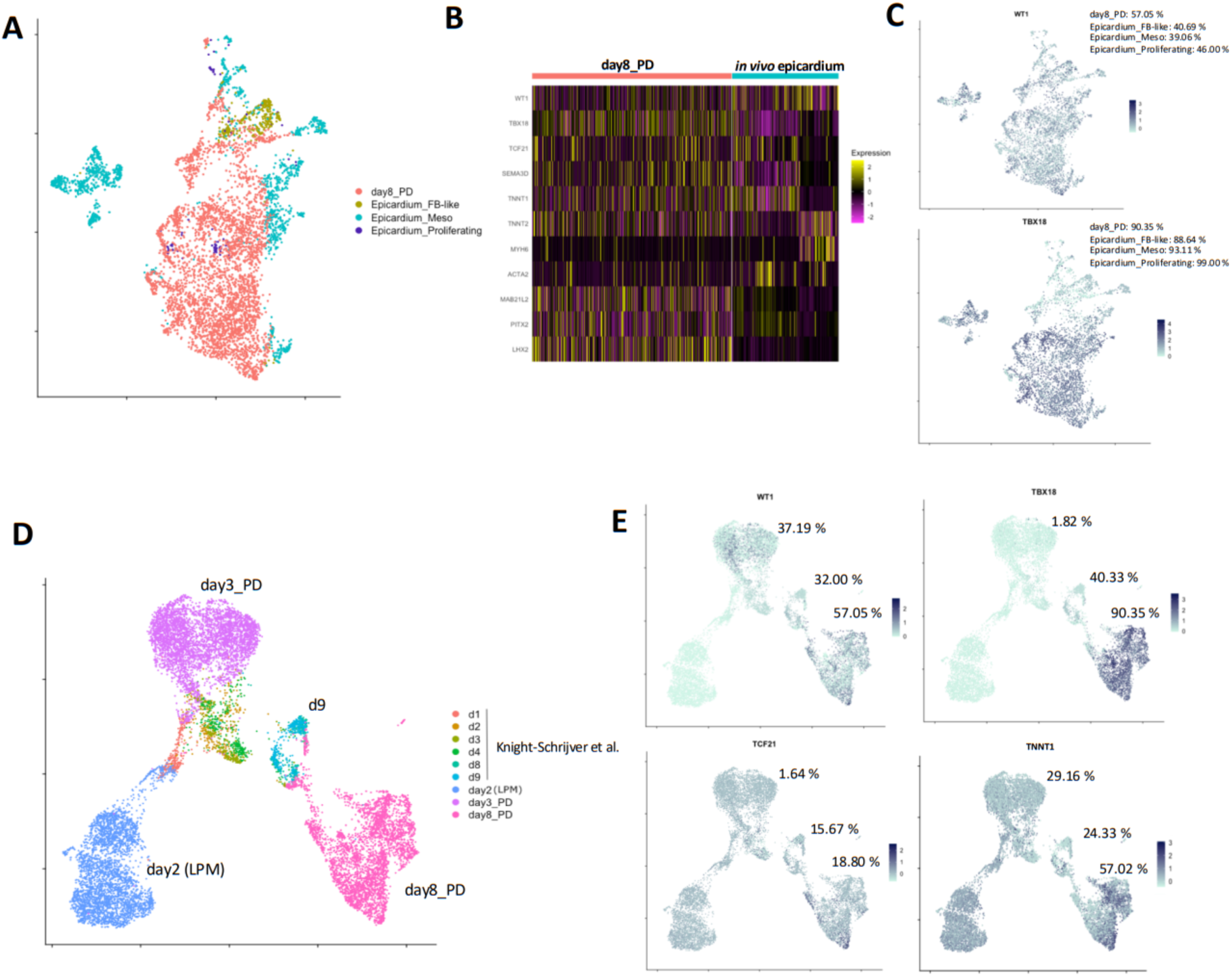
Comparison of PD-treated cells with the published scRNAseq data of the epicardium. (**A**) Integrated UMAP plot of the scRNAseq data of day 8 PD-treated cells and fetal epicardium (Epicardium_FB-like, Epicardium_Meso and Epicardium_Proliferating) [38]. (**B**) Heatmap of epicardial, pharyngeal and cardiomyocyte genes. *In vitro* cells show higher pharyngeal gene expression levels than the fetal epicardium, possibly reflecting a less differentiated state. (**C**) UMAP plots showing that *in vivo* and *in vitro* populations express similar levels of the epicardial markers *WT1* and *TBX18*. (**D**) Integrated UMAP plot of the day 2, 3 and 8 PD-treated cells with *in vitro*-differentiated epicardial cells using a published protocol [25, 38]. Cell populations from the two distinct protocols show minimal overlaps of cells representing similar differentiation stages. (**E**) UMAP plots showing that epicardial genes are enriched in the day 8 PD-treated cells. The indicated percentage refers to the fraction of the cells whose normalized transcript count is higher than 1.5, which corresponds to 0.41 on the shown natural log-transformed scale. The percentages are indicated for the samples “day3_PD”, “d9” and “day8_PD”. FB, fibroblast; Meso, mesothelial.

We also compared the gene expression profiles of PD-treated cells to the *in vitro* epicardial cells derived from human ESCs employing a published protocol (hereafter referred to as BWR) (Fig. 5D). In the BWR protocol, LPM cells were treated with BMP4, WNT3A and retinoic acid for 10 days [25], during which the authors performed scRNAseq at six different time points [38].

We integrated the data from our day 2 (LPM), day 3 and day 8 PD-treated cells with the published data from the abovementioned protocol [38] (Fig. 5E, S4). As expected, UMAP plot showed that our cells positioned closely with the BWR cells of the corresponding differentiation stages. However, the cell populations overlapped to a very limited extent, suggesting distinct cellular identities. Notably, day 8 PD-treated cells exhibited enrichment of the epicardial markers *WT1, TBX18* and *TNNT1*, suggesting that our cells could better represent *in vivo* fetal epicardium compared to the BWR cells.

In summary, the day 8 PD-treated cells expressed epicardial marker genes at levels comparable to *in vivo* fetal epicardium, and higher than previously described *in vitro*-derived epicardial cells, advocating for their use in disease and developmental modelling.

## Discussion

The research in cardiac cell derivation from human PSCs has seen a rapid advancement in recent years, as it has been emphasized not only in disease modelling, but also in drug screening and stem cell-based therapies [27, 73, 78–82]. Moreover, the induction of specific cardiac cell types, such as OFT or atrial cardiomyocytes, has been made possible by understanding signalling cues operating during cardiogenesis [83, 84]. The importance of FGF signalling has been highlighted in many contexts [16–18], but its role during early cardiac specification events still remains elusive. Here, we demonstrated that transient inhibition of the FGF-ERK signalling in LPM resulted in increased gene expression levels of proepicardial markers at the cost of FHF markers, suggesting a cell fate switch. Our cardiac cells with transient PD treatment showed expression levels of (pro)epicardial marker genes comparable to *in vivo* fetal epicardial cells and higher levels of those markers compared to the previously described and sequenced *in vitro* epicardial population [25, 38], suggesting a robust representation of *in vivo* (pro)epicardium.

However, further analyses such as electrophysiological profiling and functional assays should be performed to confirm the cell types, in order to consolidate the findings from our analyses. Since PSC-derived cardiac cultures tend to contain heterogeneous cell types [57, 85], it would also be important to test the functional integrity of the cell populations.

Recently, a cardiac field corresponding to the origin of the proepicardium has been characterized in mouse embryos and termed JCF [40–42]. It has been reported that the JCF stems from the embryonic/extraembryonic border region during gastrulation and shares a common *Hand1*^+^ progenitor population with the extraembryonic mesoderm [42]. Due to the difference in the origin of human and mouse extraembryonic mesoderm [86], the relationship of human JCF with other early progenitors remains elusive.

There is limited information on the role of ERK pulses in cardiac lineage specification. Nevertheless, it is reasonable to speculate that ERK pulses (or the lack thereof) influence the cell fates in mesodermal progenitors since they receive ERK pulses in a temporally and spatially regulated manner [16]. It has been reported that FGF signalling prevented proepicardium formation in human cells, whereas the treatment with PD173074, an inhibitor of FGF receptors, showed no difference to the control [29], consistent with our data. However, it is unclear if the timing of FGF administration corresponded to the LPM stage in this study. The FGF signalling has also been reported to play a crucial role in SHF and OFT development [87]. Looking at later developmental stages, the role of FGF for (pro)epicardial maturation has been established in chicken embryos [88]. Once the epicardium has spread on the surface of the myocardium, FGFs secreted by the myocardium promote epithelial-to-mesenchymal transition of the epicardium, inducing further differentiation [29, 89]. Conversely, FGFs secreted by epicardium have been shown to be crucial for proper myocardial proliferation and maturation [90–92]. However, substantial differences have been reported between chicken and mouse proepicardium/STM concerning the response to FGF [93], questioning the relevance of the results from chicken embryos for the mammalian systems.

On the contrary, the role of the WNT signalling for (pro)epicardium formation has been confirmed in several human *in vitro* studies [25–29]. These findings, together with our data, suggest a crosstalk between the WNT and FGF-MEK-ERK signalling pathway. Indeed, several synergistic interactions have been reported between these pathways at different levels, e.g. via β-catenin that is released and activated by ERK or via the activation of the transcription factor AP-1 or TCF4 [94, 95]. Furthermore, in zebrafish, Fgf has been reported to activate Wnt by inhibiting its antagonists, while the Wnt signalling promotes Erk activity via Gsk3 phosphorylation [96]. The abovementioned epicardial protocols typically involve a WNT inhibition phase before activating the pathway again [26–28]. Thus, inhibition of WNT might have played a similar role to the MEK/ERK inhibition described in this study. However, the timing and duration of WNT modulation vary between the protocols, necessitating further studies to elucidate the optimal conditions for proepicardium induction.

The epicardium has caught a considerable amount of attention in regenerative medicine due to its supportive role in cardiomyocyte growth during development [14, 97, 98]. Epicardial fibroblasts, dormant in the adult, reactivate upon injury, causing fibrosis and scar formation [14, 98–100]. In recent years, attempts have been made to utilize the regenerative potential of epicardium for damaged myocardium while avoiding excessive fibrosis [14, 97, 98]. In mammals, co-transplantation of ESC-derived epicardial cells and cardiomyocytes has been shown to improve cardiac proliferation, maturation and function after injury [101]. Later, epicardium-derived fibronectin has been identified as a component responsible for myocardial maturation [102]. Additionally, treatment with the epicardium-derived FSTL1 protein has been reported to stimulate cell cycle re-entry of cardiomyocytes in mouse and swine infarction models [103]. Importantly, given that the heart of zebrafish and neonatal mammals harbors a high regenerative potential, they are being used to elucidate the regenerative mechanisms [104, 105]. Single-cell RNAseq analysis of zebrafish epicardium after injury has identified a transiently activated progenitor cell population contributing to heart regeneration [106]. In the human, considerable differences have been found in the composition of fetal and adult epicardial cell populations, with a shift of gene signature from angiogenesis to immune response [38].

Thus, in light of these findings that fetal activated epicardial progenitors could promote heart regeneration, immature populations like our *in vitro*-generated proepicardial cells could be potentially applicable to regenerative therapies.

Taken together, our results implicate that in the human cell culture model, MEK/ERK inhibition at the LPM stage alters the gene expression profiles from FHF to a broad field of progenitors, including proepicardium, SHF and STM. Considering the role of fetal epicardium for cardiomyocyte regeneration, the cell population derived by our protocol could be of interest to therapeutic approaches. Future studies should address the potential of these cells using *in vitro* and *in vivo* approaches including myocardial infarction models.

## Methods

### Cell maintenance in monolayer culture

H9 primed pluripotent stem cells (WA09, WiCell) were maintained in mTeSR1 (STEMCELL Technologies, 85850) on 5 μg/mL Biolaminin 521 (BioLamina, LN521) at 20 % O_2_, 5 % CO_2_ and 37 °C. Cell passaging was performed every 4 – 5 days with TrypLE Select (Thermo Fisher Scientific, 12563011) and without ROCK inhibitor. The cell line was karyotyped and tested negative in routine mycoplasma screenings.

### Organoid culture

Pluripotent cells in the monolayer culture were dissociated with Accutase (Thermo Fisher Scientific, A1110501) and seeded in 48-well plates at a density of 400.000 cells/cm^2^ in mTeSR1 without coating and with ROCK inhibitor (VWR, 688000). From the next day, the medium was changed to mTeSR1 without ROCK inhibitor every day. Cell aggregates were kept at uniform sizes by gently pipetting up and down with a P1000 pipette.

### Cardiac differentiation

The cytokine concentrations used here correspond to the already described protocol [30]. In the monolayer culture, cells were seeded at a density of 80.000 – 120.000 cells/cm^2^ 4 days prior to the induction start, so that the wells were confluent 1 day prior to the start. In the organoid culture, induction was started 5 – 7 days after plating. Differentiation was performed in DMEM/F-12 GlutaMAX (Thermo Fisher Scientific, 10565018 and 31331028) medium supplemented with the cytokines as described below. Treatment with the respective media lasted for 1 day each time. Monolayer cultures were briefly washed with PBS before applying the new medium whereas organoids were gently washed with the new medium twice. Day 1 medium, or PS medium, consisted of 30 ng/mL Activin A (PeproTech, 120-14E), 40 ng/mL BMP4 (PeproTech, 120-05ET), 6 μM CHIR99021 (Axon MedChem, 1386), 20 ng/mL FGF2 (PeproTech, 100-18B) and 100 nM PIK90 (Axon MedChem, 1362). Day 2 medium, or LPM medium, consisted of 1 μM A8301 (Sigma, SML0788), 30 ng/mL BMP4 and 1 μM C59 (Axon MedChem, 2287). Day 3 medium contained 1 μM A8301, 30 ng/mL BMP4 and either 20 ng/mL FGF2 in the case of “control” or 1 μM PD0325901 (Axon MedChem, 1408) in the case of “PD”. Day 4 medium was composed of 25 ng/mL Activin A, 30 ng/mL BMP4 and 1 μM C59, and day 5 medium, or cardiac medium, contained 30 ng/mL BMP4, 1 μM XAV939 (Sigma, X3004) and 200 μg/mL 2-phospho-ascorbic acid (Sigma, 49752). Cells were kept in the cardiac medium until day 8 with a medium change every 1 – 2 days.

### RNA isolation

RNA was extracted using RNeasy kits (Qiagen, 74104 for monolayer cultures and 74004 for organoids). On day 8, monolayer cultures were briefly washed with PBS and collected by scraping in the RLT buffer. Organoids were washed with PBS, dissolved in the RLT buffer and passed 10-times through a 21G needle. All the following steps were performed according to the manufacturer’s handbook.

### RT-qPCR

For cDNA synthesis, Random Hexamers (Thermo Fisher Scientific, N8080127) were used as primers and SuperScript III Reverse Transcriptase (Thermo Fisher Scientific, 18080044) for reverse transcription. In the qPCR experiment, SYBR Green I (Roche Diagnostics, 04707516001) was used to make the reaction mixture, and the qPCR program was executed on LightCycler 480 II (Roche Diagnostics). The reaction efficiency was inferred from the standard curve. To normalize gene expression levels, housekeeping genes *ACTB* and *GAPDH* were used. The primers used are listed in Supplementary Table 1.

### Western blot

After washing briefly with cold PBS, cells were scraped and collected in Laemmli buffer. Sonication was performed at 20 % amplitude using Branson 450 Digital Sonifier (Marshall Scientific) 10 s at a time until a homogenous protein solution was obtained. Overheating of samples was prevented by keeping them on ice. Samples were then centrifuged at 10.000 rpm for 10 min and the supernatant was used to measure the protein concentration by NanoDrop 2000 (Thermo Fisher Scientific). 20 – 30 μg of the protein solution was adjusted to the volume of 30 μL with Laemmli buffer and boiled together with 2 μL bromophenol blue solution and 3 μL of 1 M DTT for 5 min at 90 °C. Samples were loaded in NuPAGE 4 – 12 % gels (Thermo Fisher Scientific, NP0321BOX) and separated in NuPAGE SDS running buffer (Thermo Fisher Scientific, NP0002) for 70 min at 150 V. Afterwards, the proteins on the gel were transferred to Amersham Protran nitrocellulose membrane (GE Healthcare, GE10600003) in NuPAGE transfer buffer (Thermo Fisher Scientific, NP0006-1) for 90 min at 110 V at 4 °C. The membrane was incubated in PBS with 0.1 % Tween-20 (PBSTw) for 10 min and blocked in PBSTw with 5 % skim milk powder for 1 h. Primary antibodies were dissolved in 5 % BSA in PBSTw, except for the Histone H3 antibody, which was dissolved in 1 % skim milk powder in PBSTw. The membrane was incubated with the primary antibody solution for 1 – 2 days at 4 °C. After washing the membrane 3 times for 10 min with PBSTw, the secondary antibody was applied in PBSTw with 5 % skim milk powder for 2 – 3 h. After washing the membrane 3 times for 10 min with PBSTw, the proteins were visualized using ECL Prime Western Blotting Detection Reagent (GE Healthcare, RPN2236) and imaged with ChemiDoc MP (Bio-Rad). If blotting with a different antibody was desired, the membrane was incubated in the stripping buffer (Thermo Fisher Scientific, 46430) for 15 – 60 min and the procedures were repeated from the blocking step. The antibodies used are listed in Supplementary Table 2. The intensity of the imaged bands was measured by means of the software ImageJ (v. 1.53t).

### Immunofluorescent staining of 2D cultures

Pluripotent stem cells were seeded on 8-well μ-slides (Ibidi, 80826) and differentiated towards cardiac cells as described above. On day 8, after a brief wash with PBS, cells were fixed with 4 % PFA (Thermo Fisher Scientific, 28908) for 15 min. Then, cells were permeabilized in PBS with 0.1 % Triton X-100 (PBSTr) for 10 min and blocked with CAS-Block (Thermo Fisher Scientific, 008120) for 15 min. Primary antibodies were dissolved in CAS-Block and applied on cells overnight at 4 °C. After 3 quick washes and 2 long washes of 15 min in PBSTr, secondary antibodies in CAS-Block were applied for 2 – 3 h in the dark. After 3 quick washes followed by 2 long washes of 15 min in PBSTr, 1 μg/mL DAPI was applied in PBSTr. Cells were imaged with Leica TCS SP8 (Leica Microsystems) at 63x magnification and images were processed with the software Imaris (v. 9.9). The antibodies are listed in Supplementary Table 2.

### Immunofluorescent staining of organoids

Organoids on day 8 of cardiac differentiation were briefly washed with PBS, fixed in 4 % PFA for 30 min and permeabilized with 0.5 % Triton X-100 in PBS for 30 min. Blocking was performed with CAS-Block for 1 h. Organoids were treated with primary antibodies dissolved in CAS-Block for 2 days at 4 °C. After 3 washes with PBSTr for 15 min, organoids were incubated in secondary antibodies and 1 μg/mL DAPI in CAS-Block for 1 day at 4 °C in the dark. After 3 washes with PBSTr for 15 min, organoids were put in PBSTr with 1 μg/mL DAPI. For imaging, organoids were transferred to 8-well μ-slides and let stand for 45 min to sediment. Images were taken with Leica TCS SP8 at 20x magnification and processed with Imaris. The antibodies are listed in Supplementary Table 2.

### Live-cell imaging of beating cardiac cells

Beating structures on day 8 were imaged using AF6000 (Leica Microsystems) in which the cells were kept at 37 °C. 10 images were taken per second, and the images were compiled to videos that represent the actual beating speed.

### Bulk RNA sequencing

The libraries for RNA samples extracted from monolayers and organoids were prepared and sequenced separately. 500 ng RNA were used as an input in monolayer-derived samples, whereas 120 ng were used for organoids. We prepared the libraries according to the manual for NEBNext Ultra II RNA Library Prep Kit (New England BioLabs, E7770). PCR amplification was performed for 5 cycles for monolayer-derived samples and 9 cycles for organoids. Samples were purified with AMPure XP beads (Beckman Coulter, A63881). For pooling, the cDNA quality was evaluated with Fragment Analyzer (Agilent Technologies) and the concentration was measured with Qubit Fluorometer (Thermo Fisher Scientific). Sequencing was executed on NextSeq 500 Sequencer (Illumina) at reNEW/CPR Genomics Platform (University of Copenhagen, Denmark), and the raw data were processed according to the standardized pipeline. The count matrix was analyzed using the software R (v. 4.2) [107], in which the normalization and differential gene expression analysis were performed with the package DESeq2 (v. 1.34.0) [108]. For the downstream analyses, only genes with more than 300 normalized transcript counts in 2 or more samples were retained.

### Single-cell RNA sequencing

On day 2, 3 and 8, control and PD-treated cultures were dissociated with TrypLE Select and cryopreserved in KnockOut Serum Replacement (Thermo Fisher Scientific, 10828028) supplemented with 10 % DMSO. Afterwards, samples were shipped to and handled by Single Cell Discoveries (Utrecht, The Netherlands). Cell thawing was performed using DMEM/F-12 medium (Thermo Fisher Scientific, 21331020) supplemented with 10 % KnockOut Serum Replacement. The libraries were prepared using the 10x Genomics 3’ v3.1 kit targeting 5000 cells/sample, and sequenced on NovaSeq 6000 (Illumina) targeting 80.000 reads/cell in PE150 mode. Raw data were handled by the abovementioned company. The CellRanger (v. 7.0.1) [109] output was filtered for empty barcodes and analyzed using the Seurat package (v. 4.3.0) [110] on R. Cells were filtered using the argument nFeature_RNA > 2750 & nFeature_RNA < 12000 & percent.mt < 12, retaining 17582 cells. Data from 5 samples from our study were compiled using the merge function without the integration step. The data were further normalized with LogNormalize method and scale factor 10000. The FindVariableFeatures function was set to retain 2000 features with the selection method “vst”. Cell cycle genes were regressed out in the ScaleData function. 40 dimensions (dims = 1:40) were used to create the UMAP plots. To integrate datasets from other studies, the NormalizeData function was used with LogNormalize and scale factor 10000. 2000 features, found by the FindVariableFeatures function, were used in the SelectIntegrationFeatures and FindIntegrationAnchors steps and for the integration. The integrated datasets were further processed as described above, analogously to the unintegrated dataset.

## Supporting information

SupplementaryVideo1_2D_control1

SupplementaryVideo2_2D_control2

SupplementaryVideo5_3D_control1

SupplementaryVideo6_3D_control2

## Data availability

Bulk RNAseq and scRNAseq data from this study can be found in ArrayExpress under E-MTAB-12573 and E-MTAB-12619, respectively. Furthermore, the following data from Gene Expression Omnibus were used for combined analysis: GSE216019 and GSE216177 [38].

## Statistics

P-values for qPCR experiments were calculated with Student’s 2-sided t-test using t.test() function in R. P-values in bulk RNAseq and scRNAseq experiments were determined using the respective R packages. Significance levels were defined as follows: non-significant (ns) P > 0.05, * P ≤ 0.05, ** P ≤ 0.01, *** P ≤ 0.001, **** P ≤ 0.0001. A biological replicate is an independent experiment, in which the cells were seeded and differentiated at the same time.

## Acknowledgments

We thank the staff of the DanStem/reNEW Core Facilities, especially M. Michaut, H. Wollmann, J.M. Bulkescher, A. Shrestha, M. Paulsen and E. Fernandez-Rebollo. We are grateful to the A. Grapin-Botton group for sharing the H9 cell line.

