## Supplementary data for "ERK activity as a key player in determining cardiac cell fate choices"

### Supplementary Materials

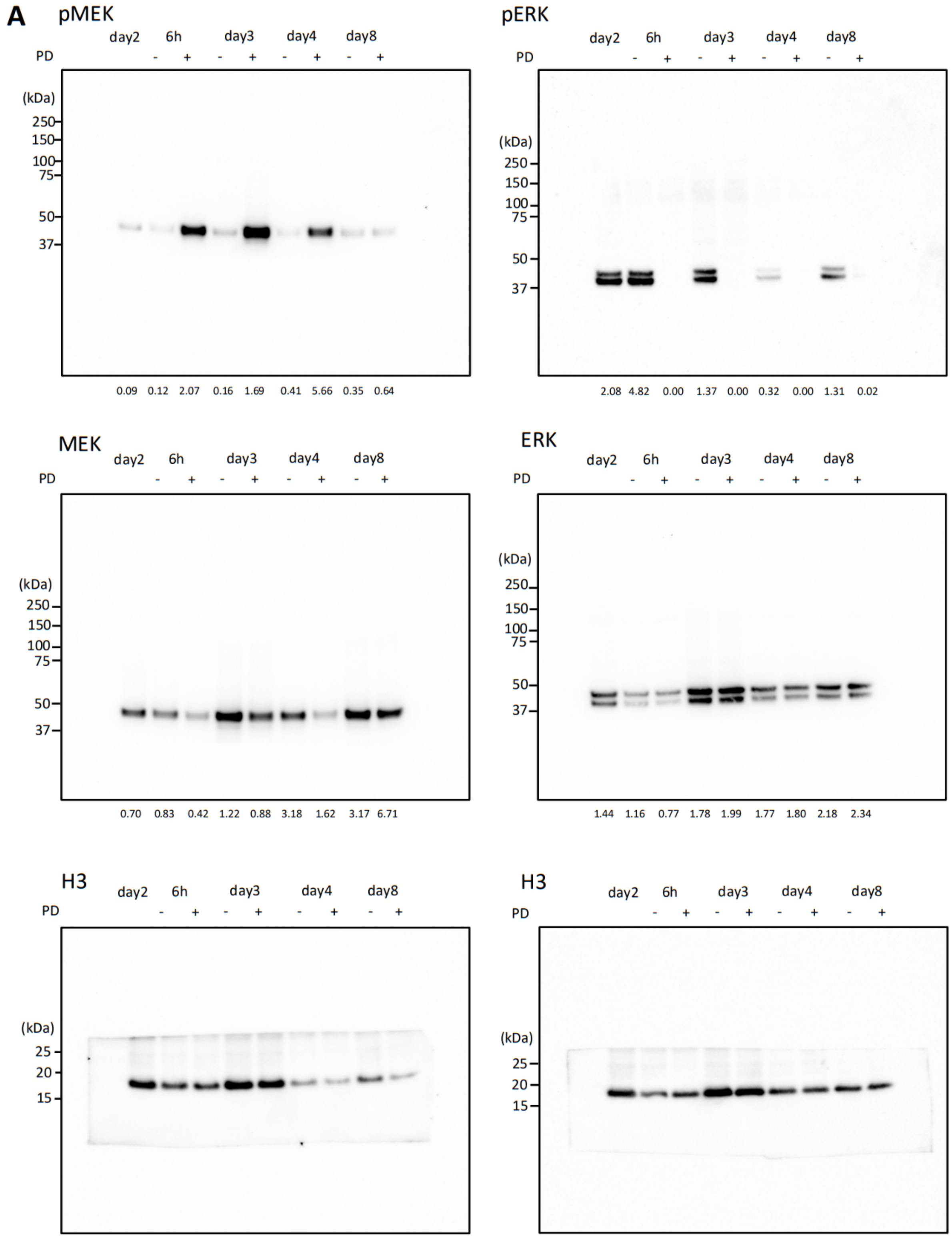


**Supplementary Figure 1**: Uncropped western blot images of biological replicates. Related to Figure 2B. The figure continues to the next page.

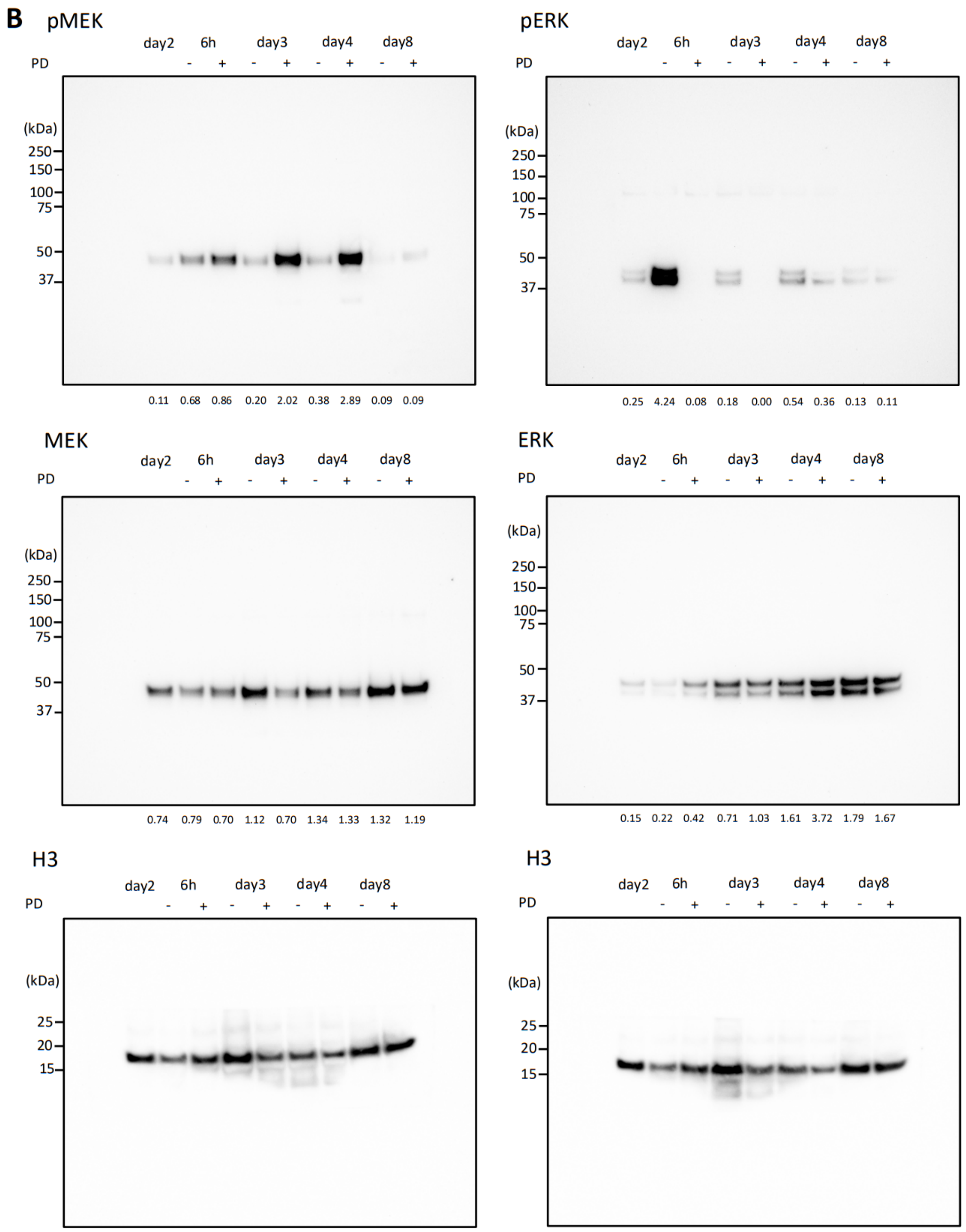


**Supplementary Figure 1.** Uncropped western blot images of biological replicates. Related to Figure 2B. Blots are shown for replicate 1 (**A**) and replicate 2 (**B**). The numbers under the blot indicate the relative intensity of the bands normalized against the Histone H3 band intensity. After protein transfer, the membrane was cut into two parts. The upper part was blotted for pMEK (pMEK1/2 = 45 kDa) or pERK (pERK1 = 44 kDa, pERK2 = 42 kDa), stripped, and re-blotted for MEK (MEK1/2 = 45 kDa) or ERK (ERK1 = 44 kDa, ERK2 = 42 kDa), respectively. The lower half was blotted for Histone H3 (15 kDa).

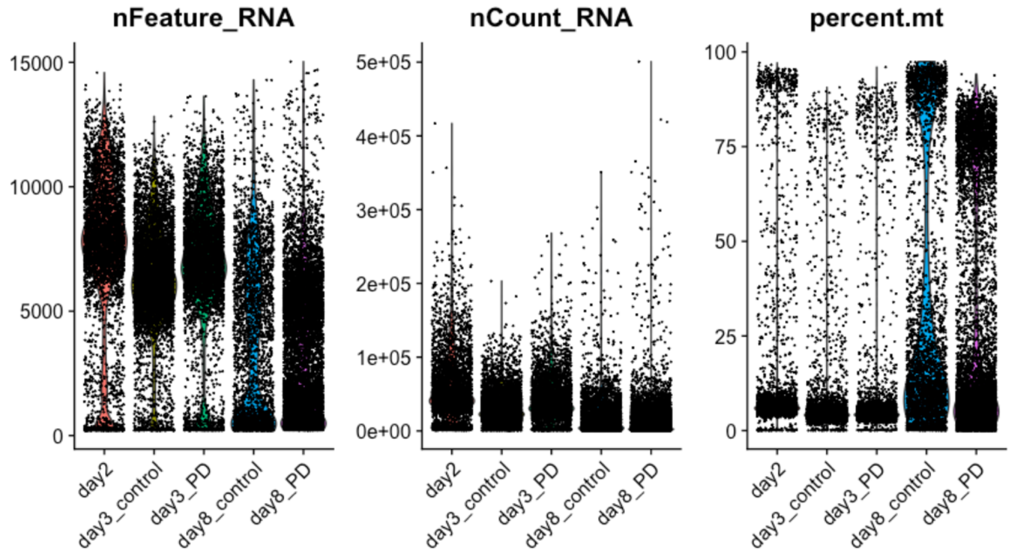


**Supplementary Figure 2.** Violin plots showing unfiltered data for nFeature_RNA (number of genes in each cell), nCount_RNA (number of transcript molecules in each cell) and percent.mt (percentage of mitochondrial genes in each cell) before the removal of low-quality cells in the scRNAseq experiment. Related to Figure 4.

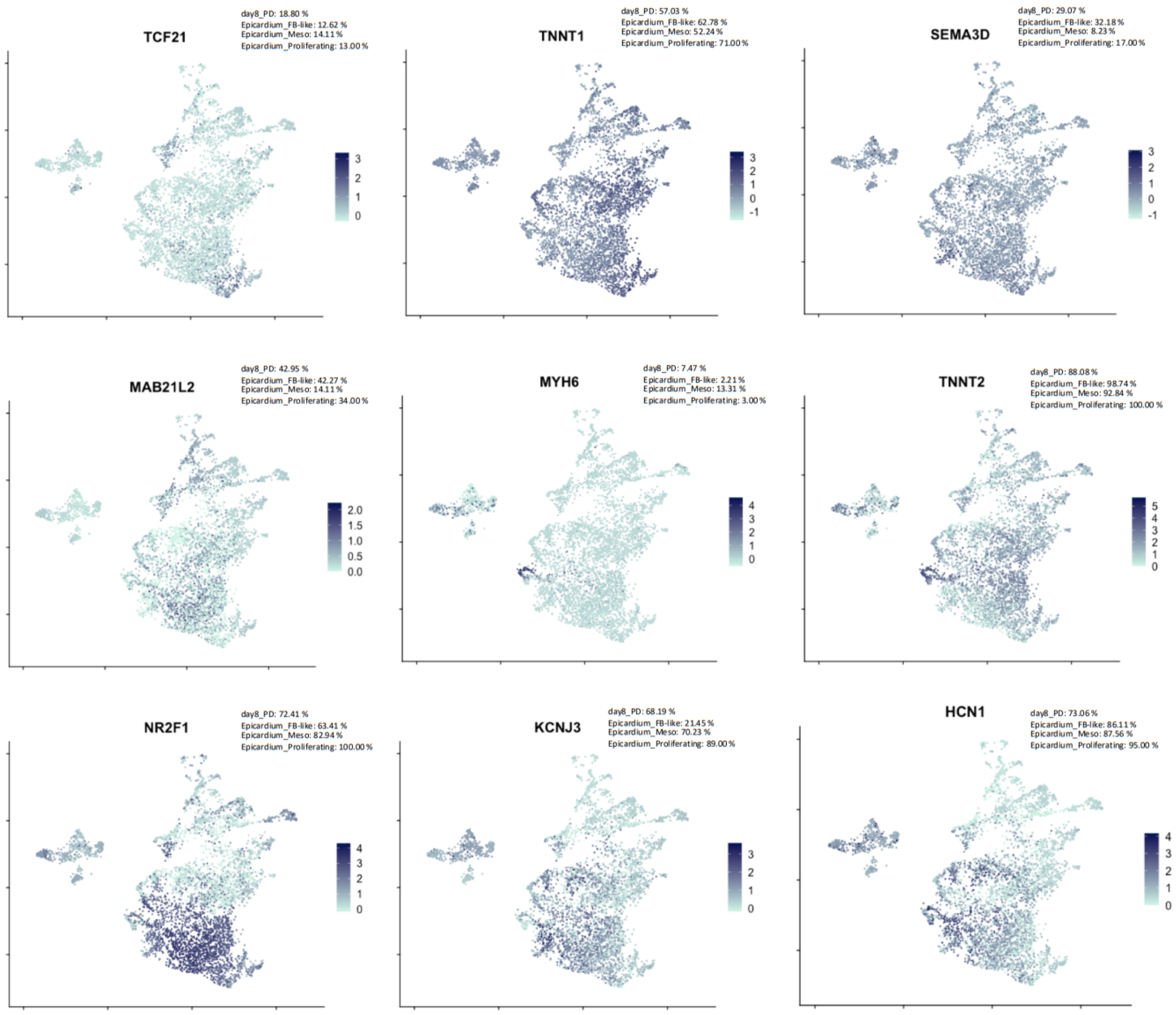


**Supplementary Figure 3.** UMAP plots showing the expression levels of cardiogenesis-related genes in day 8 PD-treated cells and fetal epicardium [38]. Related to Figure 5C. The percentage refers to the fraction of cells with the normalized transcript counts higher than 1.5, corresponding to 0.41 on the shown natural log-transformed scale.

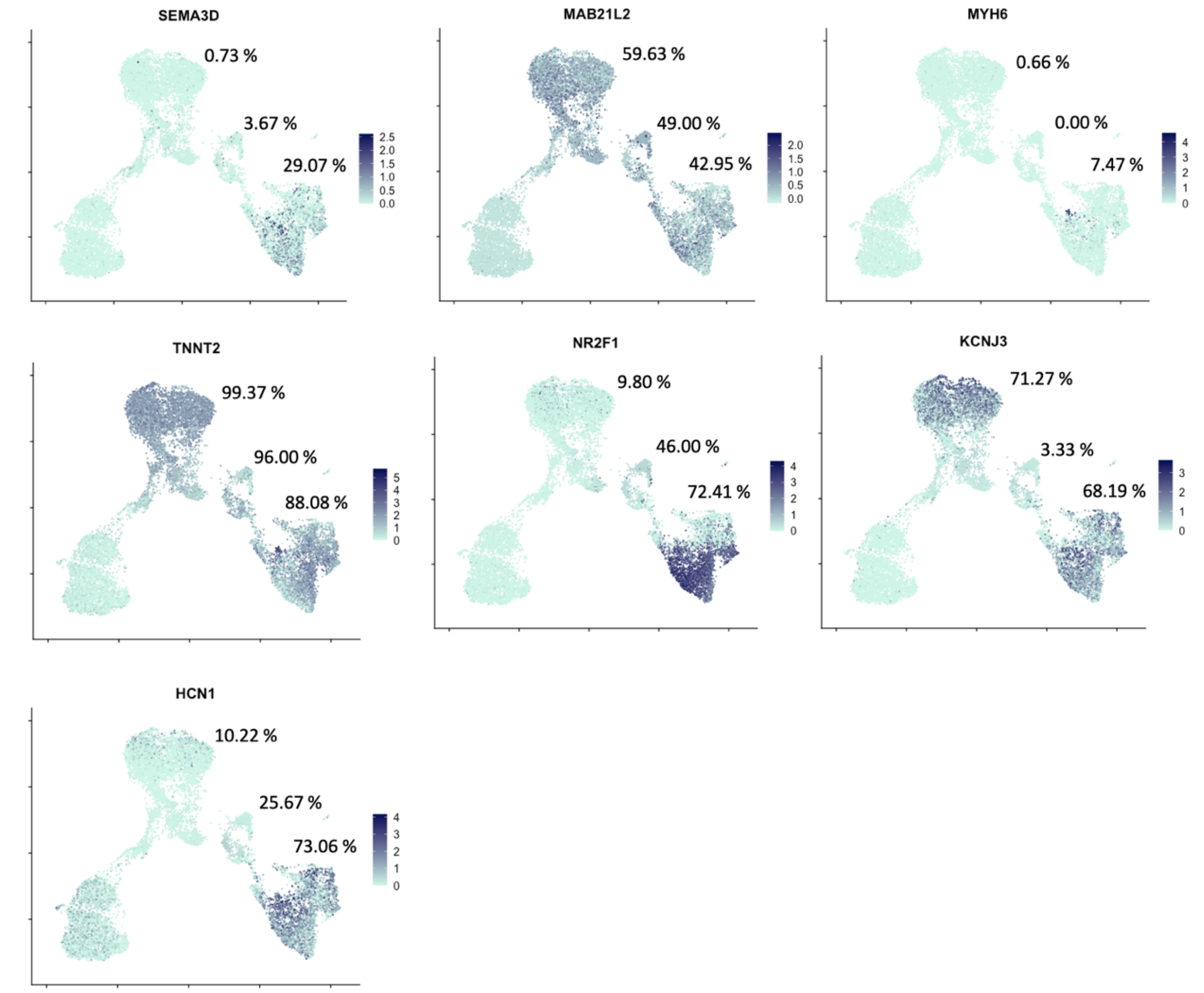


**Supplementary Figure 4.** UMAP plots comparing the cardiogenesis-related gene expression levels of PD-treated cells from this study and cells from the BWR protocol [25, 38]. Related to Figure 5E. The percentage refers to the fraction of cells with the normalized transcript counts higher than 1.5, corresponding to 0.41 on the shown natural log-transformed scale. The percentages are shown for the samples “day3_PD”, “d9” and “day8_PD”.

| Gene | Forward | Reverse |
|---|---|---|
| ACTB | CCAACCGCGAGAAGATGA | CCAGAGGCGTACAGGGATAG |
| GAPDH | AGCCACATCGCTCAGACAC | GCCCAATACGACCAAATCC |
| NKX2.5 | ACCCAGCCAAGGACCCTAGA | AGCTCATAGACCTGCGCCTG |
| TNNT1 | AGGCAGAACAGAAGCGTGGT | CAGGAGGGCTGTGATGGAGG |
| TNNT2 | TGACATAGAAGAGGTGGTGGAA | TCGTCCTCTCTCCAGTCCTC |
| WT1 | ACCAGTGTGACTTCAAGGACTGT | GCTGAAGGGCTTTTCACCTGT |

**Supplementary Table 1: List of primers**

List of primary antibodies

| **Target** | **Host** | **Assay** | **Dilution** | **Manufacturer** | **Reference no.** |
|---|---|---|---|---|---|
| Phospho-ERK | Rabbit | WB | 1:1000 | Cell Signaling | 4370 |
| Total ERK | Rabbit | WB | 1:1000 | Cell Signaling | 4695 |
| Phospho-MEK | Rabbit | WB | 1:1000 | Cell Signaling | 9154 |
| Total MEK | Rabbit | WB | 1:1000 | Cell Signaling | 8727 |
| Histone H3 | Mouse | WB | 1:5000 | Abcam | ab10799 |
| ACTA2 | Mouse | IF | 1:500 | Thermo Fisher Scientific | 14-9760-82 |
| TNNT2 | Mouse | IF | 1:200 | Thermo Fisher Scientific | MA5-12960 |
| TBX18 | Rabbit | IF | 1:200 | Abcam | ab115262 |
| WT1 | Rabbit | IF | 1:100 | Abcam | ab89901 |

List of secondary antibodies

| **Name** | **Assay** | **Dilution** | **Manufacturer** | **Reference no.** |
|---|---|---|---|---|
| Anti-Mouse HRP | WB | 1:10.000 | Thermo Fisher Scientific | A16011 |
| Anti-Rabbit HRP | WB | 1:10.000 | Thermo Fisher Scientific | A16023 |
| Anti-Mouse IgG Alexa Fluor 488 | IF | 1:200 | Thermo Fisher Scientific | A-21202 |
| Anti-Rabbit IgG Alexa Fluor 594 | IF | 1:200 | Thermo Fisher Scientific | A-21207 |

**Supplementary Table 2: List of antibodies**
